# Behind the scene of body mass allometries

**DOI:** 10.1101/2022.09.13.507740

**Authors:** Lars Witting

**Affiliations:** Greenland Institute of Natural Resources, Box 570, DK-3900 Nuuk, Greenland

**Keywords:** Allometry, Life history evolution, interference competition, bird, mammal, abundance

## Abstract

I use data based life history models for 9,488 species of birds and 4,865 species of mammals to illustrate natural selection causes for the evolution of inter-specific body mass allometries. Each model integrates the growth and demography of individuals with the life history energetics and population ecology of the species. I show *i*) how the primary selection of resource handling and mass-specific metabolism generates the net energy of individuals, *ii*) how the selected net energy generates a population dynamic feedback selection where intra-specific interactive competition selects body masses that scale in proportion with net energy on the timescale of natural selection, *iii*) how the primary selection of metabolism selects an allometric curvature where the residual mass-specific metabolism—relative to the expectation of the mass-rescaling allometry—is an initially declining function of mass in terrestrial placentals and birds, but not in marsupials and bats, *iv*) how the selection of body mass buffers ecological variation in survival, and *v*) how the joint selection of mass and optimal foraging selects the exponents of body mass allometries from the dominant spatial dimensionality of the foraging ecology.

## 1 Introduction

Allometry—the study of biological scaling—examines how the phenotype of organisms changes in proportion with changes in body mass on log scale (Snell 1892; Thompson 1917; Huxley 1932; Kleiber 1932). Mass (*w*) is seen as the independent variable with the remaining phenotypic traits (*x*) following as an allometric function

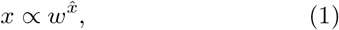

where the exponent (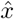, also referred to as the power) is the slope on double logarithmic scale 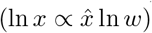.

A phenotypic scaling with mass makes intuitive sense. For organisms with about similar shapes, length tends to scale to the 1/3 power of mass, and area to the 2/3 power, reflecting the dimensional relationship between length, area, and volume. This made Rubner (1883) propose a 2/3 exponent for the allometric relation between total metabolism and mass, as expected from a thermal homeostasis that is regulated by the ratio of surface area to volume.

But an extra spatial dimension was needed (Blum 1977) to predict the observed exponents that were closer to 3/4 than 2/3 (Kleiber 1932). A fractal-like physiological network was proposed in 1999 to endow life with the missing fourth dimension, with a metabolic 3/4 power scaling suggested to follow from the constraints on optimal transports across surface areas of physiological resource transportation networks (West et al. 1999).

Even though biological traits scale with size, nothing in biology makes sense except in the light of evolution (Dobzhansky 1973). The true independent variable is natural selection most often; not body mass. The allometry point of view *is not* by itself a cause-effect relation.

Why, e.g., should metabolic scaling and other life history allometries follow from physiological constraints and a change in mass, when natural selection operates on the intra-specific variation across the joint life history of all traits? Physiological transportation networks are naturally selected to match the naturally selected metabolic rate and mass. It is the natural selection balance between metabolism and mass that defines their allometric relation (Witting 2017a,b), and this balance is not really constrained because many species with similar masses have quite different metabolic rates.

To understand the evolution of body mass allometries it is not sufficient to explain an allometric exponent from some apparent constraint on physiological variation. We need instead to build a theory that links the natural selection of body mass to the natural selection of the life history and foraging ecology as a whole; including traits like metabolism, home range size, competitive resource foraging, reproduction, survival, and lifespan. The observed inter-specific allometries should follow from the life histories and ecological traits that are predicted to follow from the selected inter-specific variation in mass.

Malthusian relativity (Witting 1997, 2008) is such a theory. It uses a single model of population dynamic feedback selection to show how the selection of metabolism defines the pace of the interactive foraging that generates net energy for population growth and them associated selection of mass (Witting 2017a,b). This predicts life histories, lifeforms, and body mass allometries in mobile organisms from replicating molecules to sexually reproducing multicellular animals (Witting 1995, 2002a, 2017a,b).

Where the original work use mathematical equations to deduce the numerical values of the allometric exponents, I use data plots to analyse the inter-specific variation in the life histories of birds and mammals, illustrating some of the essential mechanisms in population dynamic feedback selection. I show correlations that are related to the selection of body mass, metabolism, and allometries; starting with an examination of the inter-specific variation in net energy, with a separate section on the importance of metabolism for the generation of net energy. I examine the lack of evidence for *r/k*-selection, illustrate the presence of an invariant selection attractor of interactive competition, and discuss how the population dynamic feedback selection of the attractor buffers the natural selection of mass against the ecological variation in mortality.

My next focus is on the mass-rescaling selection that follows from the selection of mass, a process that selects for a decline in mass-specific metabolism and an associated dilation of natural selection time (Witting 2017a). The selected time dilation is the essential mechanism that maintains the population dynamic growth and interactive competition that is necessary to select net energy into mass during the population dynamic feedback selection of mass. I discuss how this selection of mass is entangled with the selection of optimal foraging, illustrating how the invariant selection attractor of interactive competition selects the dominant spatial dimensionality of the feeding ecology into the exponents of body mass allometries. At the end I compare the empirical and theoretical body mass allometries discussing, among others, a relation between inter-specific competition and animal abundance.

## 2 Methods

Witting (2021a) used 56,214 data on life history parameters in birds and mammals to estimate equilibrium life history models with zero population growth for 9,488 species of birds and 4,865 species of mammals. I use these models in my analysis, with the relevant parameters on individual growth, demography, life history energetics, and population ecology listed in Table 1. The details and equations for the population dynamic feedback selection of the traits are published elsewhere (Witting 1997, 2008, 2017a,b).

**Table 1:**
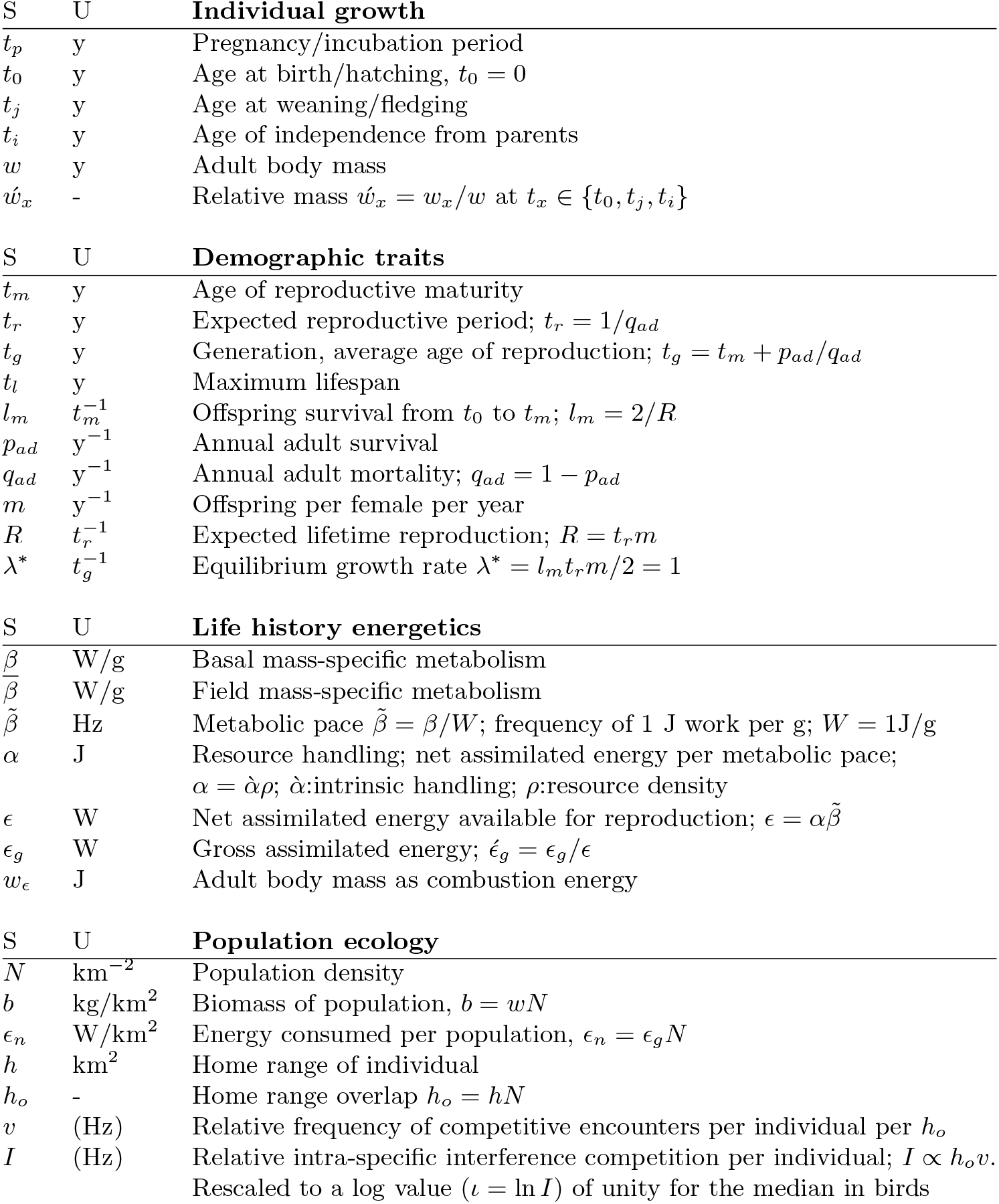
Symbols (S) and units (U) of estimated traits, with details in Witting (2021).

Several of the life history and ecological traits that are included in my analysis are not usually provided as data. Dependent upon traits and the availability of data, the trait estimates for a given species may reflect a combination of raw data, inter-specific extrapolations by allometric correlations at the lowest taxonomic level with data, and derived traits that are calculated from a combination of traits that are given either by raw data or inter-specific extrapolation (see Witting 2021 for details).

I use a double logarithmic scale to estimate traits correlations and inter-specific allometric exponents by linear regression. I differentiate between the traits that are estimated from data and inter-specific extrapolations, using only data estimates, and derived traits that are calculated from data estimates of other traits, in statistical analyses. Plots are presented with parameter estimates for all species.

Owing to major life history differences I split mammals into placentals (minus bats), marsupials and bats, while all birds are analyses together. Statistical correlations are calculated mainly for birds and placentals, as there are often too few data for marsupials and bats. The dominant spatial dimensionality of the foraging ecology follows the data based classification in Witting (2017a). All birds, bats, and marsupials are classified as having 2D ecology, while few taxa of placentals are classified with 3D ecology [Cetaceans, Primates, and the three Carnivora families Otariidae, Odobenidae, and Phocidae]. A selected set of inter-specific traits correlations are plotted in Fig. 1 for birds and placentals, and in Fig. 2 for marsupials and bats. In the sections below I refer collectively to plots of the same parameters (marked by identical letters) across the four taxonomic groups in the two figures.

**Figure 1:**
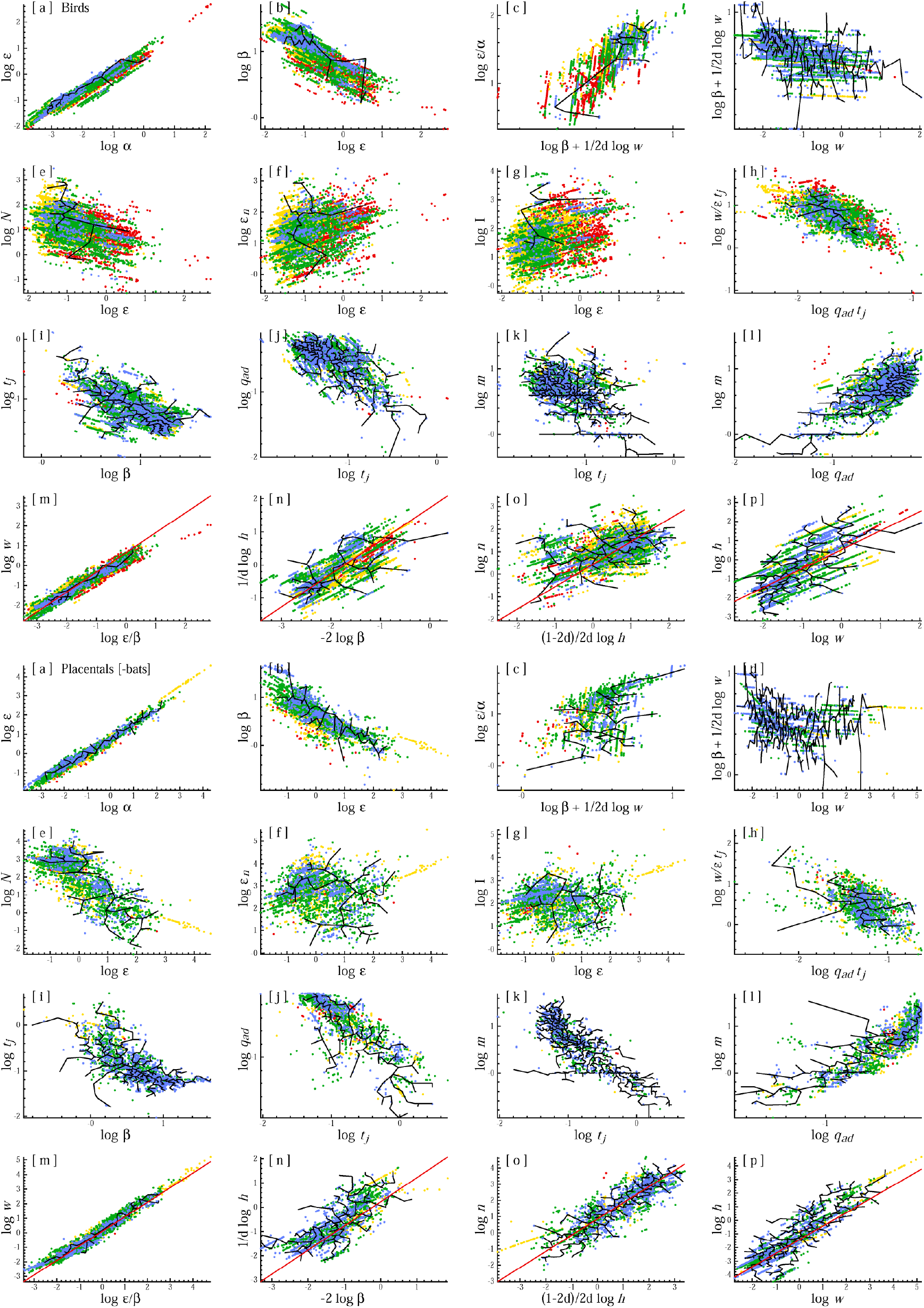
Life history evolution. Inter-specific trait correlations that illustrate the natural selection of life histories in birds (top 15 plots) and placentals (minus bats; bottom 15 plots). The black lines outline the space with data for both parameters, and the coloured dots estimates at different estimator levels, as defined by the highest estimation level for the two parameters. The red lines in plots m to p are theoretical predictions of proportionality. Estimator levels: data (black), genus (blue), family (green), order (yellow), and class (red); with points of the latter sitting on top of the former.

**Figure 2:**
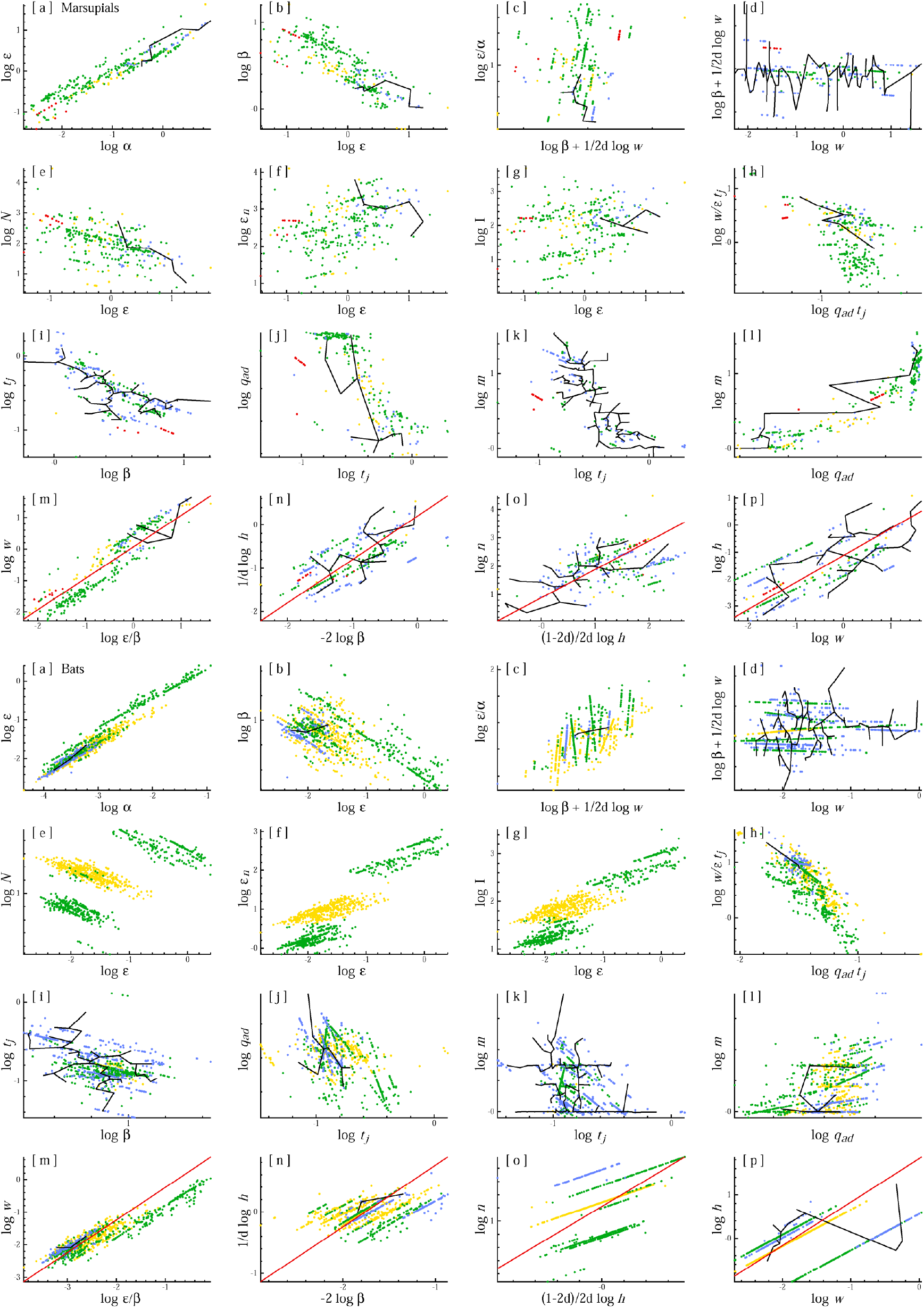
Life history evolution. Inter-specific trait correlations that illustrate the natural selection of life histories in marsupials (top 15 plots) and bats (bottom 15 plots). The black lines outline the space with data for both parameters, and the coloured dots estimates at different estimator levels, as defined by the highest estimation level for the two parameters. The red lines in plots m to p are theoretical predictions of proportionality. Estimator levels: data (black), genus (blue), family (green), order (yellow), and class (red); with points of the latter sitting on top of the former.

## 3 Net energy

The net energy (*E*) that an organism has available for reproduction is one of the most fundamental and essential traits in natural selection. It is exposed not only to primary selection because of the direct coupling to fitness through the rate of replication, but it is also the primary driver of population dynamic feedback selection. It generates the necessary energy for population dynamic growth and thus for a population density that is so large that individuals meet in interactive competition, with the frequency of the interactive encounters per individual being the main driver for the natural selection of energy consuming life history traits like body mass, multicellularity, competitive quality like interactive behaviour, and sexual and eusocial reproduction (Witting 1997, 2002a, 2008, 2017a,b).

Net energy 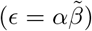 evolves from a positive primary selection on resource handling (*α*) and the pace of resource handling 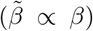, with the latter defined by mass-specific metabolism (*β*; Witting 2017a). The evolutionary scaling of metabolism, however, depends also on a mass-rescaling selection decline in response to a selection increase in mass, with the latter following from the selected increase in net energy. The final scaling between metabolism and mass depends on the relative importance of the primary selected metabolism for the evolutionary variation in net energy across the species that are compared (Witting 2017a).

If we have two taxa like ectotherms and endotherms—that differ in the primary selected metabolism and have only minor primary selected metabolism differences within each taxon—the difference in the primary selected metabolism between the two taxa is contained in the different intercepts of the metabolic allometries, while the slopes of the metabolic allometries reflect the intra-taxa variation in body mass that is selected from the intra-taxa variation in resource handling, including variation in the exploited resources (see Fig. 3 in Witting 2017b for details).

**Figure 3:**
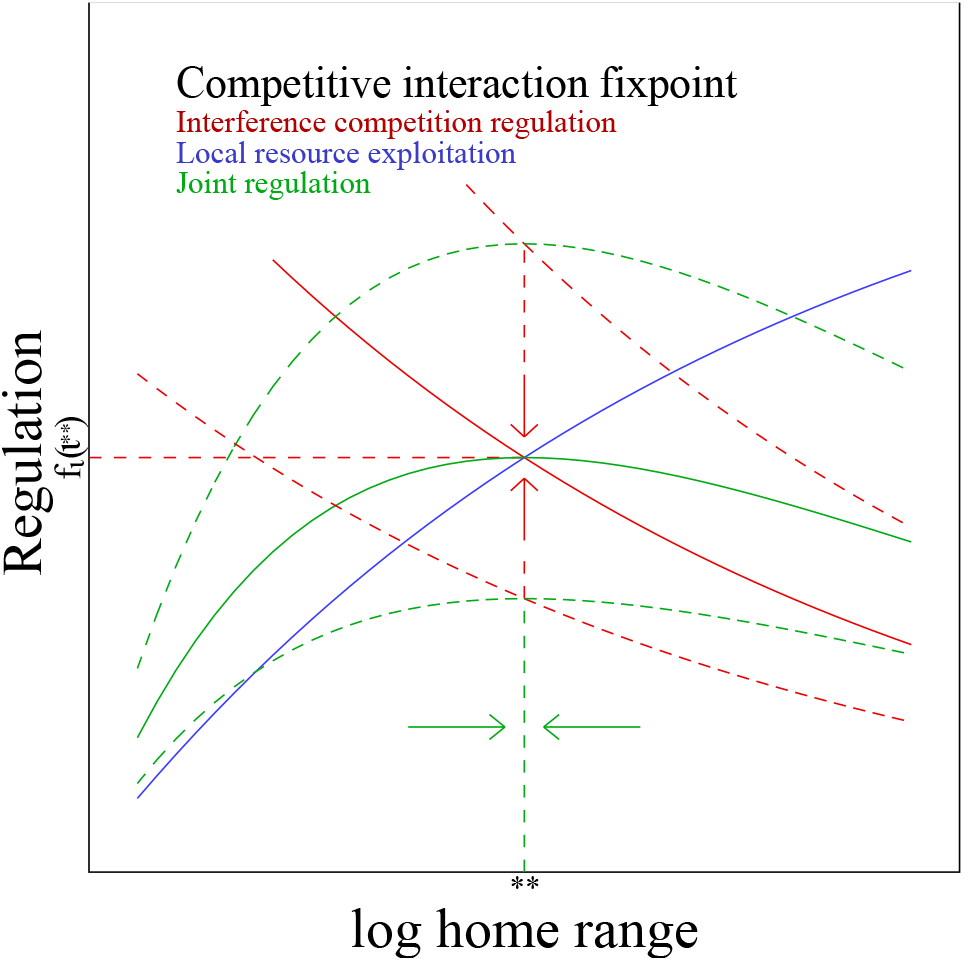
Optimal foraging. The home range of optimal density regulation (**) is defined by the selection attractor on the joint regulation (*f*_*s*_*f*_*ι*_; green curves) of local resource exploitation (*f*_*s*_; blue curve) and interactive competition (*f*_*ι*_; red curves), with the latter [*f*_*ι*_(*ι*^*∗∗*^)] being selected to match the interactive competition (*ι*^*∗∗*^) for the selection attractor on body mass. From Witting (2017b).

For birds and mammals that have approximate Kleiber (1932) scaling with *β* exponents around −1/2*d* (*d* = 2 or *d* = 3), it follows that the primary cause for the evolutionary variation in net energy within each taxon is variation in resource handling. Hence, net energy is strongly dependent on resource handling as illustrated by the a-plots [exponent: 0.64 (se:0.03; n:37) for birds & 0.73 (se:0.01; n:68) for placentals]. Mass-specific metabolism is then declining with net energy because of the secondary downscaling from a body mass that increases with net energy [b-plots, exponent: −0.36 (se:0.05; n:32) for birds & −0.32 (se:0.02; n:64) for placentals].

Some fraction of the residual variation (*β/w*^−1/2*d*^) in mass-specific metabolism (*β*) relative to Kleiber scaling (*w*^−1/2*d*^), however, should reflect differences in the primary selected metabolism. The residual net energy *E/α* that is left over from the variation in resource handling should thus be positively dependent on the residual variation in *β*; as illustrated in the c-plots [exponent: 2.34 (se:0.38; n:32) for birds & 1.03 (se:0.29; n:64) for placentals].

## 4 Primary selected metabolism

An invariant primary selection of metabolism will generate an exponential increase in mass-specific metabolism on the per-generation timescale of natural selection (Witting 2020). Yet, metabolic evolution in physical time accelerates more in the smaller species as these evolve over a larger number of generations. Witting (2018) found that this body mass dependent evolutionary acceleration was bending the metabolic allometry over time, with the primary selection of metabolism predicting a curvature in the metabolic allometry of placentals, as documented by Hayssen and Lacy (1985), Dodds et al. (2001), Packard and Birchard (2008), Kolokotrones et al. (2010), and MacKay (2011). The metabolic scaling for marsupials, on the other hand, showed almost no curvature suggesting that placentals have more primary selected variation in mass-specific metabolism than marsupials. From the stronger acceleration of primary selected metabolism in the smaller species we expect a decline in the Kleiber scaling residual for mass-specific metabolism (*β/w*^−1/2*d*^) with an increase in mass, as illustrated in the d-plots not only for placentals but also for birds. With correlation coefficients of −0.21 (p*<*0.001; n:531) and −0.46 (p*<*0.001; n:356), and regression exponents of −0.03 (se:0.01; n:531) and −0.08 (se:0.01; n:356) for placentals and birds both relations are highly significant.

The largest species, however, have the smallest mass-specific metabolism, and they may thus experience a more unconstrained selection of metabolism, with a larger per-generation increase, than the smaller species. This would explain not only the upward bend in the metabolic allometry for the largest placentals (Witting 2018), but also the non-linearity of the residual distributions in the d-plots for placentals and birds. For placentals, we have a clear decline in residual mass-specific metabolism from the smallest to medium sized species, with an apparent increase in the larger species. The pattern in birds is less clear, yet there seems to be a slight non-linearity where an initial decline is levelling off in the larger species.

In agreement with Witting (2018), a decline in the primary selected metabolism with an increase in mass could not be detected for marsupials, and nor for bats [correlation: −0.02 (p:0.84; n:74) for marsupials & 0.16 (p:0.11; n:98) for bats].

## 5 *r/k*-selection

Consider the possibility that the use of energy is controlled by frequency-independent *r/k*-selection. Owing to the frequency-independence and the associated constancy of the relative fitness of a variant, *r/k*-selection has the special feature that the average fitness of a population is a trait that evolves just like any other phenotypic trait. This implies a constant increase in the average fitness of populations, as expressed by an increase in *r* or *k* (Fisher 1930).

A selection increase in net energy by *r/k*-selection is thus associated with an increase in the demographic traits and density of a population, with an even stronger increase in the amounts of resources that are consumed by the population. There is, however, no allometric support for this very basic prediction. Population density is a declining function of net energy [e-plots, exponent: −0.78 (se:0.35; n:26) for birds & −1.21 (se:0.13; n:65) for placentals], and the consumption of energy by populations are invariant of net energy in birds, but not in placentals where population consumption declines with increased net energy [f-plots, correlation: −0.06 (p:0.78; n:26) for birds & −0.23 (p:0.07; n:65) for placentals].

Another issue with *r/k*-selection relates to the quality-quantity trade-off, where a few large or many small offspring can be produced from the same amount of energy (Smith and Fretwell 1974; Stearns 1992). This implies a replication rate that is inversely related to mass, with continued selection for a decline in mass. The existence of large organisms is simply not supported by basic *r/k*-selection, making it essential to identify another selection mechanism that will outbalance the frequency-independent selection for negligible mass.

## 6 Competitive interaction fixpoints

The selection that follows from the density-dependent interactive competition among the individuals in a population was shown by Witting (1997) to generate a density-frequency-dependent feedback selection that outbalances the frequency-independent selection for the absence of mass. This allows for the evolution of a variety of body masses that reflect the underlying inter-specific variation in the net assimilated energy of species, with *r* and *k* being selected—as observed—to decline with a selection increase in net energy and body mass (Witting 1997, 2000, 2002b).

This population dynamic feedback selection is actively balancing the density-frequency-dependent selection for an increase in mass against the frequency-independent selection for a decline in mass. It is the frequency-independent selection for a decline in mass and increase in *r* and *k* that generates population dynamic growth with the level of interactive competition increasing with an increase in abundance. The increase in interference competition is then generating a densityfrequency-dependent resource bias in favour of the competitively superior individuals, and when the bias becomes sufficiently strong it outbalances the selection of the quality-quantity trade-off selecting net energy into mass at the cost of the demographic traits.

The attractor of population dynamic feedback selection is a competitive interaction fixpoint that maintains an invariant level of intra-specific interference by the selection allocation of net energy between reproduction and mass. By selecting the level of interference, the attractor is exactly outbalancing the negative selection of the quality-quantity trade-off by adjusting the resource bias of interactive competition and the associated selection of mass.

The log of the average number of interactive encounters per individual (*ι*) at the competitive interaction fixpoint has a predicted value of

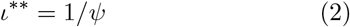

when the body mass in selected to an evolutionary equilibrium (∗∗), and a value of

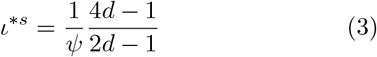

when body mass is selected to increase exponentially at an evolutionary steady state [Witting, 1997; *ψ* is the intra-population gradient (around the average life history) in the fitness cost of interference competition per unit interference (caused by differential resource access); ∗*s* denotes steady state; *d* is the dominant number of spatial dimensions in the foraging ecology of the species].

The truly invariant component of the attractors is not the number of interactive encounters, but the intrapopulation fitness bias from interactive competition [*ι*^*∗∗*^*ψ* = 1 & *ι*^*∗s*^*ψ* = (4*d −* 1)/(2*d −* 1)]. To understand population dynamic feedback selection it is essential to understand that it are the population level gradients in resource assess by interactive competition (*ι*^*∗∗*^*ψ & ι*^*∗s*^*ψ*) that are the core attractors of natural selection, with all the other life history traits being naturally selected by the feedback mechanisms of the selection so that the population mechanistic generation of intra-specific interference is adjusted to match the pre-determined value of the overall attractor.

With non-significant correlations between net energy and the estimated level of interference competition we do not falsify the existence of competitive interaction fixpoints [g-plots, correlation: 0.31 (p:0.28; n:14) for birds & −0.15 (p:0.33; n:41) for placentals].

## 7 Extrinsic mortality

The selection attractors of the competitive interaction fixpoints imply a feedback where increased extrinsic mortality selects for increased fecundity. This is imposed by the selection of invariant interference, where increased extrinsic mortality—and the associated decline in density and interference competition—selects an increase in fecundity until the interference competition of the competitive interaction fixpoint is reestablished (Witting 1997, 2008).

The energy for a selection increase in fecundity is taken primarily from a selection decline in mass. Yet a large component of the increase in yearly fecundity with increased yearly mortality follows from the body mass selected scaling of rates and periods (see next subsection). So, to analyse for the influence of extrinsic mortality, we examine the residual component where an allometrically independent increase in mortality selects for an increase in fecundity. The residual variation in body mass (*w/*ϵ*t*_*j*_)—relative to the mass expectation of the net assimilated energy 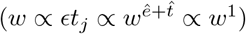— should thus be selected to decline with increased extrinsic mortality. This is illustrated in the h-plots where residual mass is declining significantly with a mortality rate (*q*_*ad*_*t*_*j*_) that is time-scaled to eliminate the allometric dependence [correlation: −0.76 (p*<*0.001; n:37) for birds & −0.57 (p*<*0.001; n:68) for placentals].

A decline in mass with an increase in natural or anthropogenic mortality has been identified in a variety of studies (e.g., Reznick et al. 1996; Haugen and Vøllestad 2001; Sinclair et al. 2002; Carlson et al. 2007; Herczeg et al. 2009; Rossetto et al. 2012). The often-observed decline in mass following human-induced mortality is often argued to reflect a selective harvest of especially the larger individuals (e.g., Browman 2000; Sinclair et al. 2002; Olsen et al. 2004). Yet, the importance of intra-specific mass-specific mortality is likely overstated in the literature, because an evolutionary decline in mass is the base-case response of natural selection to increased mortality.

## 8 Mass-rescaling selection

Metabolism implies that population dynamic feedback selection cannot select mass without an additional scaling of the life history (Witting 2017a). A potential selection increase in mass implies that the offspring metabolises more energy during the period of parental care, and this additional waste of energy needs to be compensated for in one way or the other in order to maintain the reproductive output that generates the population growth and interactive competition that selects energy into mass.

A solution to the problem is mass-rescaling selection for a decline in mass-specific metabolism as this can maintain the net energy and reproductive output on the timescale of natural selection (Witting 2017a). The mass-rescaling decline in metabolic pace with an increase in mass implies less net assimilated energy in physical time, but because life-periods scale inversely with metabolic pace, the decline in metabolism implies prolonged life-periods with net energy and reproduction being maintained on the dilated timescale of natural selection.

Mass-rescaling selection can be identified by a selection decline in mass-specific metabolism with an increase in the net energy that drives the selection of mass [b-plots, exponent: −0.36 (se:0.05; n:32) for birds & − 0.32 (se:0.02; n:64) for placentals]. This implies an inverse scaling between mass-specific metabolism and biological periods [i-plots, exponent: −0.74 (se:0.05; n:255) for birds & −0.60 (se:0.04; n:409) for placentals], a scaling that maintains the average net energy on the per-generation timescale of natural selection.

The inverse scaling of rates and periods is reflected in a decline in yearly mortality [j-plots, exponent: − 0.77 (se:0.03; n:492) for birds & −0.79 (se:0.05; n:171) for placentals] and fecundity [k-plots, exponent: −0.66 (se:0.02; n:1280) for birds & −0.98 (se:0.02; n:1273) for placentals] with increased life periods. This results in a strong positive correlation between yearly mortality and yearly fecundity [l-plots, exponent: 0.78 (se:0.03; n:512) for birds & 0.93 (se:0.04; n:165) for placentals]. Another consequence of mass-rescaling selection is an invariant relation where the energy that is selected into body mass is proportional to net energy on the 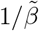 timescale of the species (Witting 2017a), i.e., body mass is selected as *w ∝* ϵ*/β* as illustrated in the m-plots [theoretical exponent of 1; observed of 0.92 (se:0.04; n:32) for birds & 1.00 (se:0.03; n:64) for placentals].

A body mass that is selected proportional to 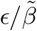 does not in itself indicate a particular scaling between metabolism and mass (or net energy and mass). The values of the allometric exponents are not given by the mechanisms considered so far.

## 9 Allometries from optimal foraging

With the selection attractor on body mass determining the level of interference competition in the population, the selection of mass is intrinsically linked to the ecology-ical regulation of the foraging process that generates the net energy that drives the population dynamic feedback selection of mass. This link is so strong that the exponents of the mass-rescaling allometries follow from the spatial constraints on optimal foraging (Witting 1995, 2017a).

Optimal foraging is a balance between the cost (i.e. regulation) of local resource exploitation and the cost of interactive competition (Fig. 3). The former selects for large home ranges to avoid self-inhibition from an over-exploitation of the local resource, and the latter selects for small home ranges to avoid competitive encounters with other individuals in the overlapping areas of individual home ranges.

For cases with a selection increase in mass we find that the reduced metabolism from mass-rescaling will generate a reduced pace of foraging, and thus also a decline in the level of interactive competition. This implies an increase in the relative cost of local resource exploitation with a selection adjustment for an increase in the size of the average home range until the level of interference is re-established at the competitive interaction fixpoint. And with regulation (*f*) by interference [*f*_*ι*_(*ι*^*∗∗*^*ψ*)] being fixed by the invariant selection attractor of interactive competition (*ι*^*∗∗*^*ψ*), regulation by local resource exploitation (*f*_*s*_) is also a selected invariance, and so is the amount of resources metabolised by the population (Witting 1995, 2017a,b).

These invariant regulation levels are the top-down constraints of the overall selection attractor. The regulations, however, are generated bottom-up by the foraging behaviour of the individuals in the population, with the natural selection solution to the latter being restricted to patterns that generate the invariant regulation of the overall selection attractor.

To examine the possible trait combinations that can generate the regulation of the overall selection attractor, we may deal with regulation as a multiplicative function on net energy (where *f* = 1 is no costs, and *f* = 0 maximal cost). Local resource exploitation 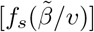 may then be approximated as a monotonically increasing function of the period (1*/v*) between reuse of foraging tracks (*v* is pace of reuse) when scaled by the metabolic pace of the organism to obtain a relative function that reflects pace at the limit with infinite home ranges (Witting 1995, 2017a). The pace of track reuse (*v* = *v*_*f*_ */l*) is foraging speed (*v*_*f*_) divided by track length (*l*), with foraging speed on the body mass axis being inversely proportional to metabolic pace 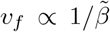 Garland 1983; Calder 1984). With track length scaling to the 1*/d*th power of the home range because of the *A* = *L*^2^ and *V* = *L*^3^ relations between length (*L*), area (*A*), and volume (*V*), foraging speed scales as 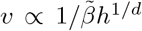 with regulation by local resource exploitation scaling as *f*_*s*_(*β*^2^*h*^1*/d*^).

As the *β*^2^*h*^1*/d*^ argument of the *f*_*s*_ function is invariant at the selected foraging optimum, we expect a *h*^1*/d*^ *∝ β*^−2^ relation where the d*th* root of the home range scale in proportion to the negative square of mass-specific metabolism [n-plots, theoretical exponent of 1; observed of 0.57 (se:0.09; n:74) for birds, 0.84 (se:0.06; n:190) for placentals, & 0.89 (se:0.23; n:23) for marsupials]. This relation does not by itself give us a value for the scaling of metabolism and home range with body mass, but it gives us a scaling between home range and mass-specific metabolism, with the relation being dependent on the dominant dimensionality of the foraging ecology.

If we look at regulation by interactive competition we have *f*_*ι*_(*h*_*o*_*v*) = *f*_*ι*_(*Nh*^(*d−*1)*/d*^*/β*), as the level of interference is approximated by the product between home range overlap (*h*_*o*_ = *Nh*) and the pace of track reuse (*v ∝* 1*/βh*^1*/d*^). Now, with *β ∝* 1*/h*^1/2*d*^ from the invariance of local resource exploitation, we may rewrite regulation by interference as *f*_*ι*_(*Nh*^(2*d−*1)/2*d*^). Thus, from the invariant argument we expect a *N ∝ h*^(1−2*d*)/2*d*^ relation where population density scales in proportion with the −3/4th and −5/6th power of the home range for two-dimensional versus three-dimensional ecological systems [o-plots, theoretical exponent of 1; observed of 0.40 (se:0.06; n:126) for birds, 0.98 (se:0.03; n:296) for placentals, & 0.63 (se:0.11; n:40) for marsupials].

The invariant amount of resource metabolised by populations (*Nwβ ∝ w*^0^) is illustrated by the 1*/Nβ ∝ w* relation in the p-plots [theoretical exponent of 1; observed of 0.83 (se:0.07; n:162) for birds, 1.18 (se:0.04; n:326) for placentals, & 0.80 (se:0.10; n:50) for marsupials]. We may thus for the invariant *Nh*^(*d−*1)*/d*^*/β* argument of interference regulation exchange *N* by 1*/wβ* and obtain *h*^(*d−*1)*/d*^*/wβ*^2^. This invariance implies proportionality between body mass and *h*^(*d−*1)*/d*^*/β*^2^, as illustrated for placentals in the second plot in Fig. 4 [theoretical exponent of 1; observed of 1.19 (se:0.07; n:74) for birds, 1.18 (se:0.03; n:190) for placentals, & 0.95 (se:0.11; n:23) for marsupials].

**Figure 4:**
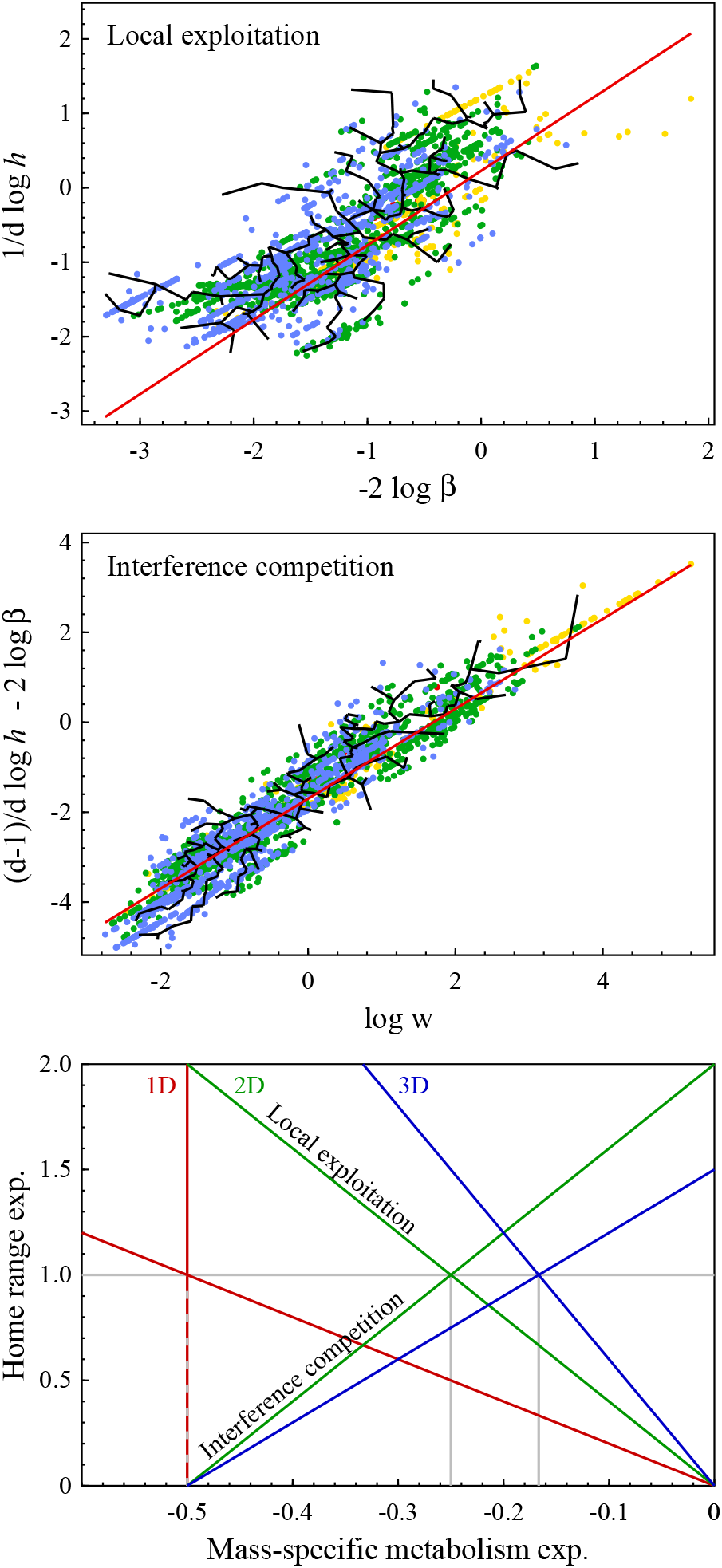
Allometric deduction. For placentals, the top and middle plots show allometric representations of the invariant regulations by local exploitation and interference competition, with the theoretical predictions (2 red lines) solved in the bottom plot for the mass-rescaling exponents 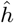 and 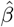 for home range and mass-specific metabolism. With 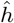 increasing with 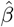 for interference competition 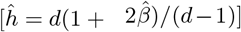 and declining for local exploitation 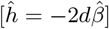, the selected exponents 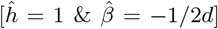 are the solutions with identical relations for *d* ∈ {1, 2, 3}.

The two proportional relations *h*^1*/d*^ *∝ β*^−2^ and *h*^(*d−*1)*/d*^*/β*^2^ *∝ w*—which reflect the invariances of the functional arguments for the regulation of foraging by local exploitation and interference competition (Fig. 4, 2 top plots)—contain the essential information that will give us the numerical values of the exponents of the body mass allometries. To see this we exchange home range and mass-specific metabolism with their mass-rescaling relations (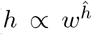 and 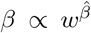), to obtain 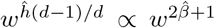 from the *h*(*d−*1)*/d/wβ*2 *∝ w*0 argument of interference regulation, and 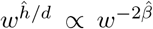 from the *β*^2^*h*^1*/d*^ *∝ w*^0^ argument local resource exploitation. This gives us the following two equations 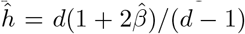 and 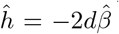 for the allometric constraints on the foraging process. These equations are solved graphically for the allometric exponents 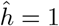 and 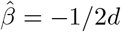 in the third plot in Fig. 4.

## 10 Body mass allometries

The final allometric exponents depend also on the relative importance of mass-specific metabolism for the primary selected net energy that drives the population dynamic feedback selection of mass (Witting 2017a). A 2D metabolic scaling with a 3/4 exponent follows when the inter-specific variation in net energy is generated from inter-specific variation in the handling and density of resources, with the exponent increasing to 7/4 when all of the inter-specific variation in net energy is generated by inter-specific variation in mass-specific metabolism. With mammals and birds having Kleiber-like scaling with typical 2D exponents of about 3/4, I focus on the mass-rescaling allometries in this section.

The estimates of allometric exponents for raw data on birds, marsupials, bats, and 2D versus 3D placentals (minus bats) are listed in Table 2. The table includes also the predicted exponents from Witting (1995, 2017a) assuming that the inter-specific body mass variation follow from inter-specific variation in resource handling and density.

**Table 2:**
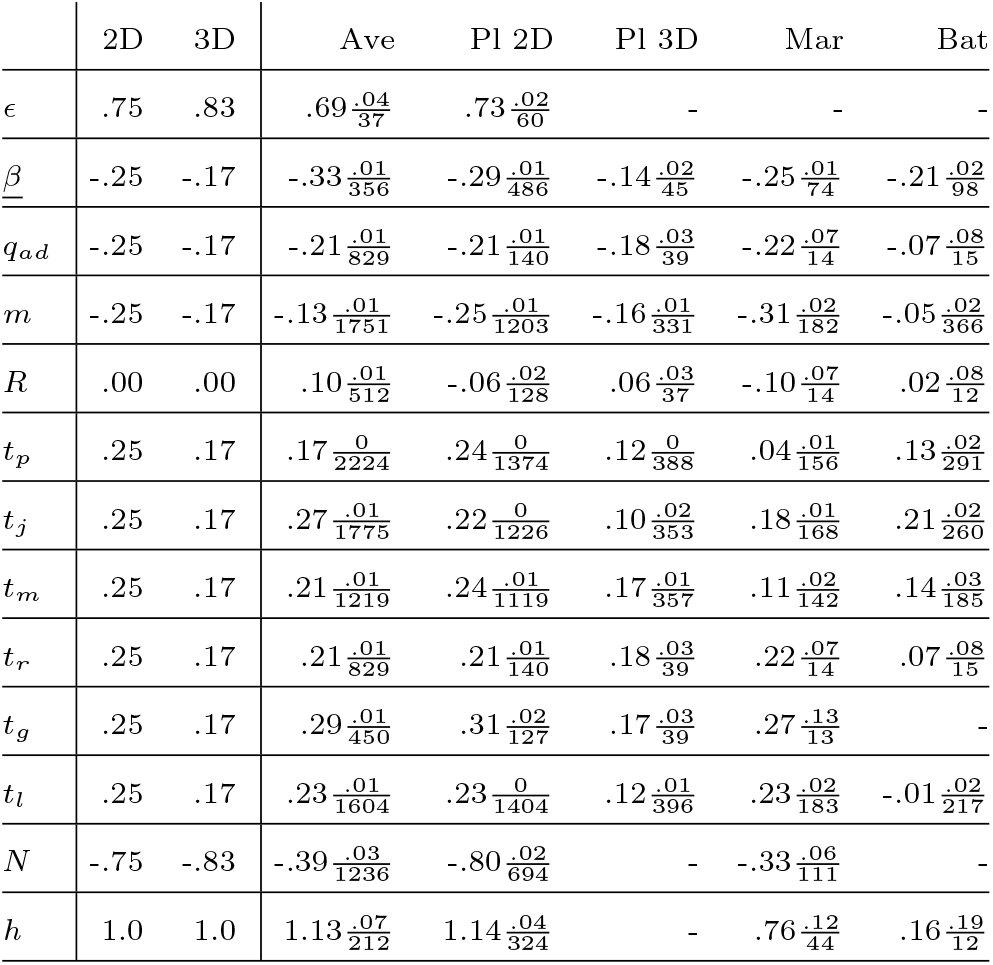
Allometric exponents estimated from data. For birds (Ave), placentals minus bats (Pl) for 2D and 3D for-aging, marsupials (Mar), and bats (Bat). The 2D and 3D columns list the theoretical exponents from Witting (1995, 2017a). Fraction numerators are se, and denominators *n*. Only *n >* 11 cases are shown.

With a linear relationship between the allometric exponents of the data and those estimated from the life history models of all 9,488 species of birds and 4,865 species of mammals (Fig. 5), we note that the allometric scaling of the data is transferred to the scaling of the estimates of missing parameters.

**Figure 5:**
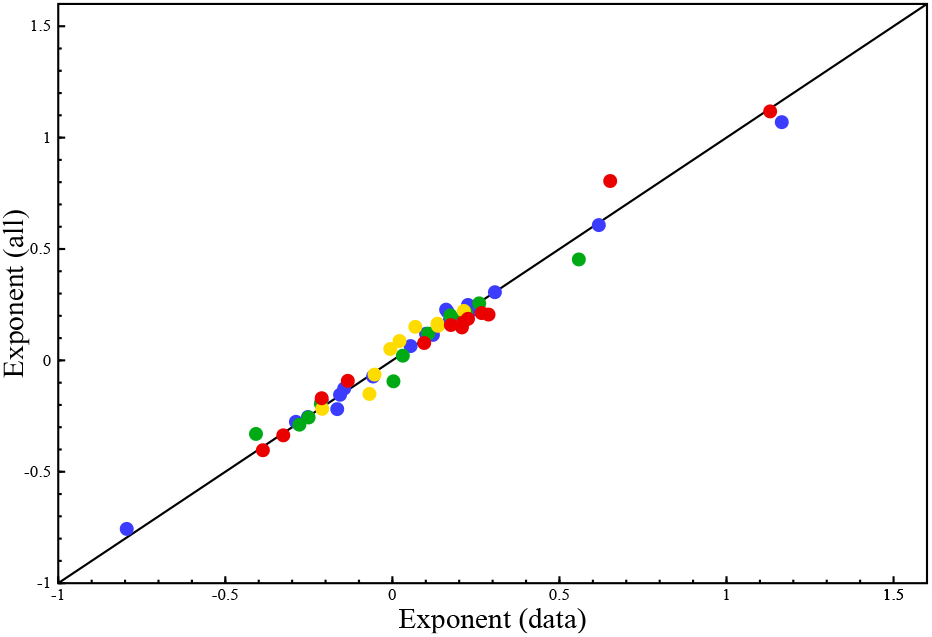
The allometric exponents from data (Table 2) compared with the corresponding exponents across all species (data plus missing parameter estimates). Red:birds; Blue:placentals; Green:marsupials; Yellow:bats.

As expected for an ecological prediction, there is a general—but not complete—resemblance between the observed and predicted exponents. A 2D-3D-like transition is marked in placentals for all life history periods and ages, as well as for metabolism, mortality, and fecundity. The exponents for birds resemble a predominately 2D-ecological competition for territories and resources.

The metabolic exponent for bats resembles the 2D expectation, however, several of the exponents for bats deviate from the predicted, including a maximum lifespan exponent of −0.01 (se:0.02) and a home range exponent of 0.16 (se:0.19). These deviations identify a potential mismatch between the interactive ecology of bats and the foraging model behind the allometric deduction. Being based on intra-specific interactions in overlapping home ranges of evenly distributed reproducing units, the foraging model was never intended to approximate the interactive ecology of gregarious bats that roots in large colonies. An extension with more elaborate models of interactive foraging is required to make predictions strictly applicable to special cases.

## 11 Abundance and inter-specific competition

The allometric deduction predicts a change in abundance 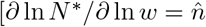 with 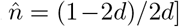 following an evolutionary change in mass. While the prediction differs from the empirical scaling for birds and marsupials in Table 2, the abundance allometry is known to change with the scale of observation (Nee et al. 1991). This change is predicted by a scale-dependent adjustment of the inter-specific variation in population dynamic feedback selection.

The allometric deduction reflects the invariant resource bias (*ι*^*∗∗*^*ψ*) across the individuals in the population. Yet the use of this invariance in the standard formulation for the allometric deduction of abundance contains the implicit assumption of an invariant resource gradient (*ψ*). This assumption seems fair on a scale where the partitioning of resources is unaffected by inter-specific competition, but it is a simplification when we examine allometries across competitive guilds. Among the species of competitive guilds we expect an inter-specific partitioning of resource by interactive competition, with the larger species having access to more resources than the smaller species. Assuming that the individuals of a species compete less per encounter when resources are abundant, this implies an inter-specific body mass dependence ln 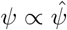 ln *w* with a 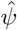-exponent that declines from zero to some lower negative value with an increase from zero in the level of inter-specific resource partitioning. Now, from the *ι ∝* 1*/ψ* dependence of eqns 2 and 3 and the expected proportionality between abundance and the number of interactive encounters, we have *N ∝* 1*/ψ*. The expected change in abundance with body mass is thus

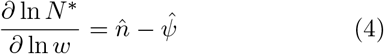

Nee et al. (1991) used the unusually good population size estimates of British and Swedish birds to examine the relationships between body mass and abundance. At the scale of all species—where we can expect a 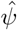 exponent around zero owing to the general lack of competition between waterfowl, shorebirds, raptors, and passerines—Nee et al. (1991) estimated a partial *∂* ln *n*^*∗*^*/∂* ln *w* relation of −0.75 among British birds, and −0.77 among Swedish, as predicted by a predominantly two-dimensional distribution of territories.

For comparisons across more closely related species, there were often positive relations between abundance and body mass in agreement with a strong partitioning of resources from inter-specific competition. Nee et al. (1991) did not quantify these relations, yet the 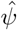 exponent needs to approach −3/4 to generate a positive relation. At the scale of the abundance data behind Table 2, we estimate a 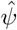 exponent of inter-specific competition of about −0.36 and −0.42 for birds and marsupials, and no apparent inter-specific resource partitioning for placentals.

